# IFN-γ stimulated murine and human neurons mount anti-parasitic defenses against the intracellular parasite *Toxoplasma gondii*

**DOI:** 10.1101/2021.11.10.468098

**Authors:** Sambamurthy Chandrasekaran, Joshua A. Kochanowsky, Emily F. Merritt, Joseph S. Lagas, Ayesha Swannigan, Anita A. Koshy

## Abstract

Dogma holds that *Toxoplasma gondii* persists in neurons because neurons cannot clear intracellular parasites, even with IFN-γ stimulation. As several recent studies questioned this idea, we used primary murine neuronal cultures from wild-type and transgenic mice in combination with IFN-γ stimulation and parental and transgenic parasites to reassess IFN-γ dependent neuronal clearance of intracellular parasites. We found that neurons respond to IFN-γ and that a subset of neurons clear intracellular parasites via immunity regulated GTPases. Whole neuron reconstructions from mice infected with parasites that trigger neuron GFP expression only after full invasion revealed that ∼40% of these *T. gondii*-invaded neurons no longer harbor parasites. Finally, IFN-γ stimulated human stem cell derived neurons showed a ∼ 50% decrease in parasite infection rate when compared to unstimulated cultures. This work highlights the capability of human and murine neurons to mount cytokine-dependent anti-*T. gondii* defense mechanisms *in vitro* and *in vivo*.

## Introduction

A select number of highly divergent intracellular microbes (e.g. measles virus, polio virus, *Toxoplasma gondii*) cause infections of the central nervous system (CNS). Though these microbes infect many cell types, in the CNS, neurons are often preferentially infected. One commonly cited reason for this neuron predominance is that neurons lack the ability to mount traditional cell-intrinsic immune responses (Joly et al., 1991; Oldstone et al., 1986; Rall et al., 1995). For example, neurons have low baseline levels of MHC I and STAT1 (Joly et al., 1991; Neumann et al., 1995; Rose et al., 2007; Wong et al., 1984). However, considerable evidence shows that neurons can respond to type I and type II interferons (Delhaye et al., 2006; Neumann et al., 1997, 1995; Rose et al., 2007) and clear certain viral pathogens (e.g. Sindbis virus (Binder and Griffin, 2001; Orvedahl et al., 2010) and vesicular stomatitis virus (Detje et al., 2009)). Limited work has been done on the capabilities of neurons to clear non-viral pathogens.

*Toxoplasma gondii* is a protozoan parasite that naturally infects most warm-blooded animals, including humans and mice. In most immune competent hosts, *T. gondii* establishes a persistent or latent infection by switching from its fast, growing lytic form (the tachyzoite) to its slow growing, encysting form (the bradyzoite). In humans and mice, the CNS is a major organ of persistence and neurons are the principal cell in which *T. gondii* cysts are found (D. J. Ferguson and Hutchison, 1987; D. J. P. Ferguson and Hutchison, 1987; Melzer et al., 2010; Cabral et al., 2016). IFN-γ is essential for control of *T. gondii* both systemically and in the CNS (Suzuki et al., 1989, 1988), in part through the activation of the immunity regulated GTPase system (IRGs) which is critical for parasite control in hematopoetic and non-hematopoetic cells (Collazo et al., 2002; Halonen et al., 2001; Taylor et al., 2000). Based upon these findings and *in vitro* studies showing that *T. gondii* readily invades murine astrocytes and neurons, but only IFN-γ- stimulated astrocytes— not IFN-γ-stimulated neurons— clear intracellular parasites (Jones et al., 1986; Halonen et al., 1998, 2001; Schluter et al., 2001), our model of CNS toxoplasmosis was that during natural infection parasites enter the CNS, invade both astrocytes and neurons, after which astrocytes kill the intracellular parasites, leaving the immunologically incompetent neuron as the host cell for the persistent, encysted form of the parasite.

Several recent findings have called this model into question. Pan-cellular ectopic expression of an MHC I allele (H-2 Ld) associated with low levels of CNS persistence (Blanchard et al., 2008; Brown et al., 1995) leads to a lower CNS parasite burden than when mice lack expression of this MHC I allele in neurons only (Salvioni et al., 2019). Moreover, the use of a Cre-based system that permanently marks CNS cells that have been injected with *T. gondii* proteins (Koshy et al., 2012, 2010), revealed that parasites extensively interact with neurons and that the majority (> 90%) of these *T. gondii*- injected neurons do not actively harbor cysts (Cabral et al., 2016; Koshy et al., 2012). Together these *in vivo* studies question our prior model by raising the possibility that neurons clear intracellular parasites.

Given these conflicting *in vitro* and *in vivo* findings, here we used primary murine neuronal cultures from wild-type and genetically modified mice in combination with cytokine stimulation and parental and transgenic parasites, including a new engineered *T. gondii*-Cre line, to reassess the ability of neurons to clear intracellular parasites in the setting of IFN-γ stimulation. These data reveal that neurons respond to IFN-γ, including up-regulating the IRGs, and that a subset of neurons (∼20%) clear intracellular parasites via the IRGs. In addition, in Cre reporter mice infected with *T. gondii*-Cre parasites that mark CNS cells only after fully invasion, whole neuron reconstructions showed that ∼40% of these *T. gondii*-invaded neurons no longer harbor parasites. Finally, IFN-γ stimulation of human stem cell derived neurons (huSC-neurons) led to an ∼50% decrease in parasite infection rate when compared to unstimulated, infected cultures. Collectively, these data highly suggest that IFN-γ stimulation leads to parasite resistance in murine and human neurons and that a subset of murine neurons clear intracellular parasites both *in vitro* and *in vivo*, likely via the IRGs.

## Results

### IFN-γ stimulated primary pure murine cortical neurons show classical IFN-γ responses

As *T. gondii* primarily infects and encysts in the cortex (Berenreiterová et al., 2011; Boillat et al., 2020; Mendez et al., 2021), we sought to determine the response of cortical neurons to IFN-γ stimulation. To accomplish this goal, we exposed pure primary murine cortical neuronal cultures to 100 U/ml of IFN-γ or vehicle control for 4 and 24hrs, followed by harvesting of total RNA. We chose these time points because prior work showed that hippocampal neurons have a delayed IFN-γ response (Rose et al., 2007). After harvesting the RNA, we used quantitative real time-PCR (qRT-PCR) to quantify the transcripts levels of traditional IFN-γ-response genes (*STAT1*, *IRF1*, *MHC-I*) as well as the effector components of the IRG system (*Irga6*, *Irgb6*, and *Gbp2*) (Howard et al., 2011; Khaminets et al., 2010). We found that *STAT1*, the classical transcription factor that drives the expression of many IFN-γ response genes, and *IRF1* were highly upregulated (4hrs:3-5 log2 fold; 24hrs: 5-7 log2 fold) compared to unstimulated neurons (**Fig 1A**), while MHC-I showed a more modest level of upregulation (∼ 2 log2 fold). In addition, consistent with finding in non-neuronal murine cells, compared to unstimulated neurons, IFN-γ stimulated neurons also significantly up-regulated *Irga6*, *Irgb6*, and *Gbp2* (4hrs:4-7 log2 folds; 24hrs: 7-9 log2 folds) (**Fig 1A**) (Boehm et al., 1998; Degrandi et al., 2013; Lafuse et al., 1995). To determine how these increased transcript levels translated to protein levels, we isolated total protein lysates from unstimulated and IFN- γ stimulated cultures. For STAT1, we blotted both for total STAT1 and for phosphorylated STAT1, the active form that enters the nucleus and binds DNA. Compared to unstimulated cultures, IFN-γ stimulation cultures showed a >10-fold increase in protein levels for total STAT1 and p-STAT1 at 24hrs post-stimulation and an >35-fold increase at 48hrs post-stimulation (**Fig 1B**). The undetectable level of total STAT1 in unstimulated neurons is consistent with previously published data suggesting that cultured neurons have low or no constitutive amounts of STAT1 (O’Donnell et al., 2015). Similarly, at 24 hours post-stimulation, the IRG complex effector proteins Irga6 and Irgb6 increased ∼ 7-fold and 10-fold respectively over unstimulated cultures, a level that was maintained at 48 hours post stimulation.

**Fig 1.**
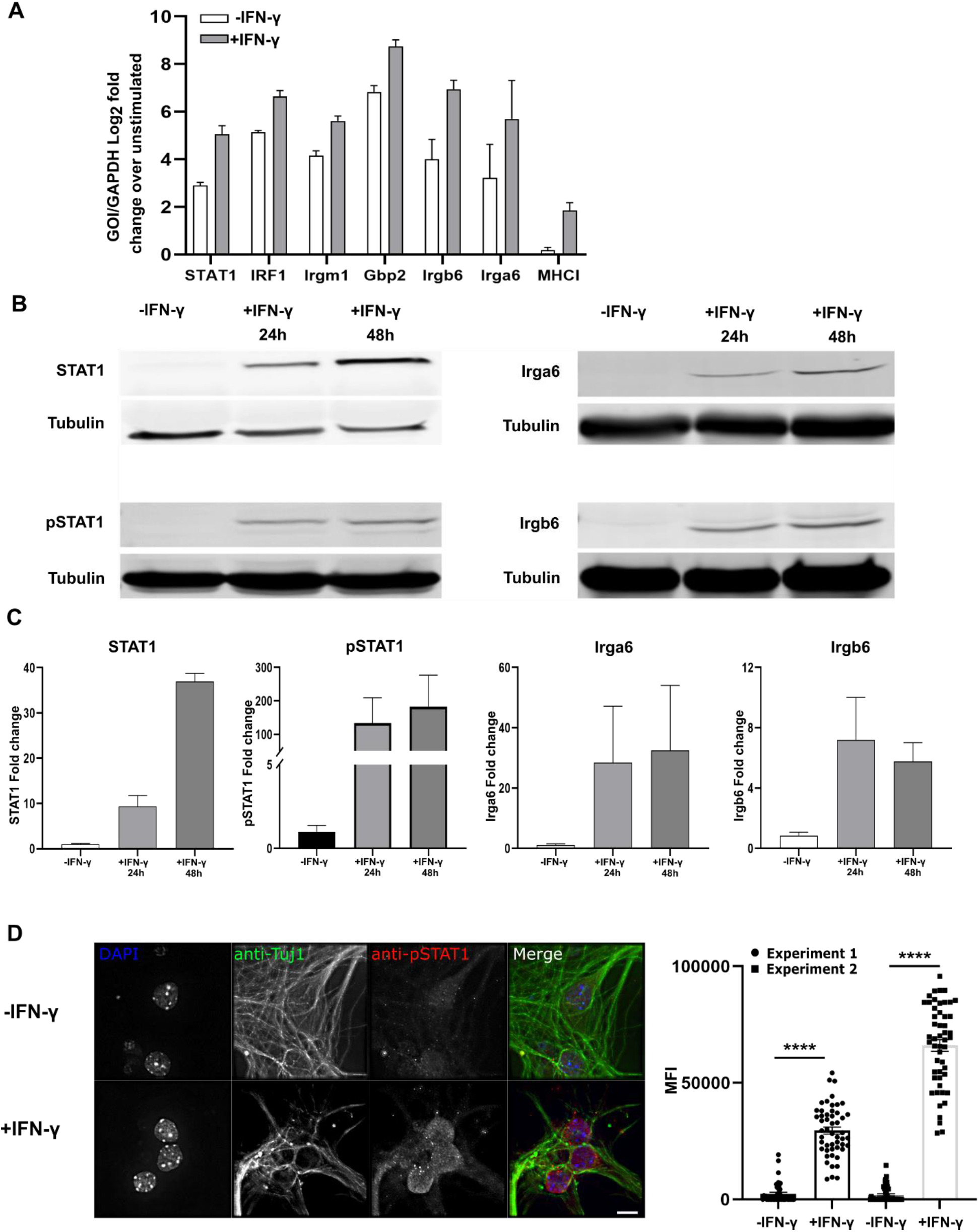
IFN-γ stimulated primary murine neurons show intact IFN-γ signaling pathway and express genes involved in *T. gondii* clearance. Primary neurons were cultured for 12 days *in vitro* (DIV), after which they were stimulated with vehicle or IFN-γ (100 U/ml). At the listed times, RNA or protein was isolated or immunofluorescent assays were performed. A. Quantification of specified genes using quantitative PCR. Expression is shown as Log2 fold change compared to unstimulated cultures. B. Representative images of western blots for specified proteins from unstimulated and IFN-γ stimulated neuron cultures. C. Densitometric quantification of western blots from (B). Densitometry of given gene is normalized to densitometry of β-tubulin and then shown as fold change compared to unstimulated cultures. For (A, C) Bars, mean ± SEM. N = 3 independent experiments. D. *Images*: Representative images of unstimulated or IFN-γ stimulated neurons stained as indicated (anti-Tuj1 antibodies stains neurons). Scale bar = 5µm. *Graph*: Quantification of the mean fluorescent intensity (MFI) of pSTAT1 nuclear signal in unstimulated or IFN-γ stimulated Tuj1^+^ cells (neurons). N = 48-51 nuclei evaluated/condition/experiment, 2 independent experiments. Bars, mean ± SD. ****p ≤ 0.0001, unpaired t-test with Welch’s correction.

To confirm that the detected changes were primarily driven by neurons and not by glial cells that commonly cause low levels of contamination in “pure” neuronal cultures, we stained the cultures to determine what percentage of cells were neurons, astrocytes, or microglia. These analyses showed that our cultures were consistently 95% neurons and 5% astrocytes; no microglia were observed (**Fig S1**). To determine the level of nuclear translocation pSTAT1 at the single neuron level, we stained unstimulated and stimulated (100 U/ml for 24hrs) primary neuron cultures with DAPI and antibodies against Tuj1, a neuronal marker, and p-STAT1. We then quantified the pSTAT1 signal intensity in Tuj1^+^ nuclei. In unstimulated cultures, we observed almost no pSTAT1 signal in Tuj1^+^ nuclei (MFI 2236 ± 353.24, mean ± SEM) (**Fig 1C, D**). Conversely, IFN-γ stimulated neurons showed robust pSTAT1 staining in Tuj1^+^ nuclei (MFI 44778 ± 25785, mean ± SEM) (**Fig 1C, D**).

Together these data show that IFN-γ stimulated primary murine cortical neuron cultures upregulate IFN-γ response genes and proteins in a delayed manner consistent with what has been observed in hippocampal neurons (Rose et al., 2007). The upregulated genes and proteins include *STAT1* and the IRG genes known to be required for IFN-γ- dependent killing of intracellular parasites in murine non-neuronal cells (Khaminets et al., 2010). Collectively, these data suggest that IFN-γ stimulated murine cortical neurons upregulate the appropriate machinery to clear intracellular parasites via the IRG system.

### IFN-γ pre-stimulation leads to a decrease in the percentage of *T. gondii*-infected neurons

To address neuronal capability for clearing intracellular parasites, we infected IFN-γ stimulated or unstimulated neurons with the two canonical parasite strains (type II and type III) that have moderate to mild acute virulence in mice because both are IRG- sensitive (Khaminets et al., 2010). We then monitored infection rates of neurons at 3, 12, and 24 hours post infection (hpi). At 3hpi, stimulated and unstimulated neurons showed a similar rate of neuron infection, regardless of infecting strain (**Fig 2A, B**). By 12 and 24 hpi, regardless of infecting strain, IFN-γ stimulated cultures showed an ∼25% decrease in neuron infection rate compared to unstimulated cultures (**Fig 2A, B**).

**Fig 2.**
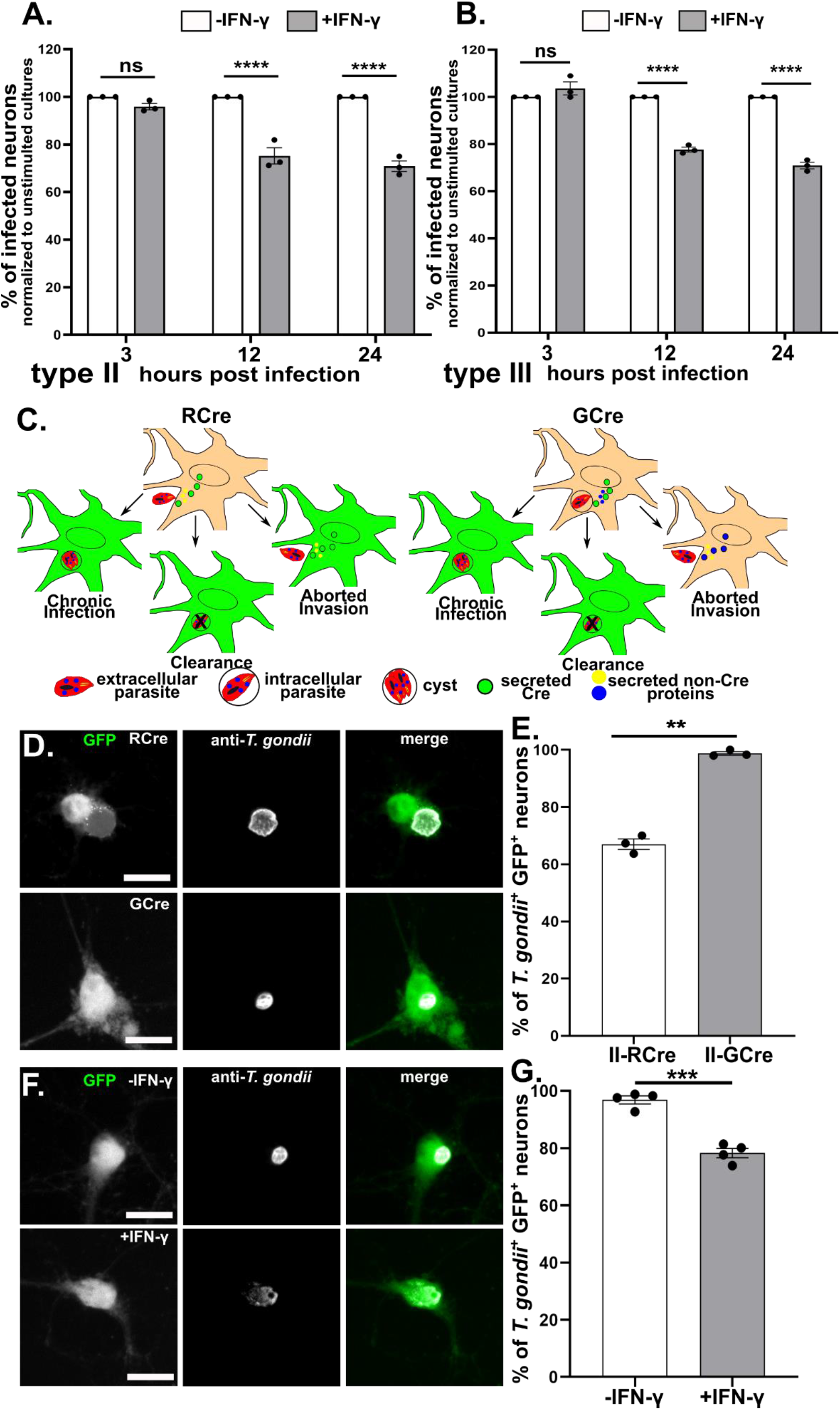
IFN-γ stimulated neurons clear intracellular parasites. Primary neurons were cultured and stimulated as in Fig 1. **A.** Graph of the percentage of infected neurons at listed time points for type II parasites, normalized to the unstimulated culture. **B.** As in (**A**) except for type III parasites. (**A, B**) Primary neurons were stimulated with IFN-γ (100U/ml) or vehicle for 24hrs, after which the cultures were infected with type II or type III parasites. At the listed time points, cultures were fixed, stained with anti-NeuN antibodies (stains neuron nuclei) and DAPI, and analyzed on an Operatta CLS high content analysis microscope. Bars mean ± SEM. N = 25-30 FOV (∼750-1000 neurons/experiment), 3 independent experiments. ****p<0.0001. ns = not significant, 2-way ANOVA with Sidak’s multiple comparisons. **C.** Schematic of host cells labeled by RCre versus GCre parasites. *Left*, In the RCre system the Cre fusion protein is secreted prior to invasion of neurons, which leads to GFP-expressing neurons that can arise from: i) injection, invasion, and chronic infection, ii) injection and invasion of host cell followed by host cell killing or clearance of the parasite, iii) injection of the protein *<u>without</u>* invasion (aborted invasion). *Right*, In the GCre system the Cre fusion protein is secreted post invasion, which leads to GFP-expressing neurons that can arise from: i) invasion and persistent infection or ii) invasion followed by killing or clearance of the parasite, but not from aborted invasion. **D.** Representative images of GFP^+^ neurons infected with RCre or GCre parasites. *Merge image*: Green = GFP-expression in neurons, white = parasites stained with anti-*T. gondii* antibodies (a cocktail of anti-SAG1 and anti-SRS9 antibodies to capture both tachyzoites and bradyzoites.) Scale bar = 10 µm **E.** Graph of the percentage of actively infected GFP^+^ neurons at 72 hpi (unstimulated cultures). **F.** Representative images of unstimulated or IFN-γ (100 U/ml) stimulated primary neurons infected with GCre parasites for 72hrs. *Merge image*, as in (**D**). Scale bar = 10 µm **G.** Graph of the percentage of actively infected GFP^+^ neurons at 72 hpi. (**E, G**) Bars mean ± SEM. N ≥ 200 GFP^+^ neurons/well, 3 wells/experiment, 3-4 independent experiments. **p ≤ 0.005, ***p ≤ 0.0005, t-test with Welch’s correction.

### GCre-expressing parasites show that neurons clear parasites in response to IFN-γ

While the prior data suggested that IFN-γ stimulated neurons might clear intracellular parasites, they could also be explained by decreased rates of late invasion in the IFN-γ stimulated neurons, especially as clearance assays in neurons are limited by the inability to synchronize infection or vigorously wash off uninvaded parasites as either procedure causes widespread neuronal death. To address the possibility of an invasion defect versus true clearance of intracellular parasites, we required a way to specifically track neurons that were infected and subsequently cleared the intracellular parasite. As noted above, we had previously developed a Cre-based system that leads to GFP expression only in host cells injected with *T. gondii* proteins. In this system, Cre is fused to a rhoptry protein (or ROP), which are parasite proteins that are injected into host cells *<u>prior to invasion</u>*, which means that parasite-triggered host cell expression does not require parasite invasion (i.e., aborted invasion) (**Fig 2C**). Thus, the RCre (ROP::Cre)-expressing parasites do not help us distinguish between aborted invasion versus clearance of intracellular parasites. Therefore, we fused Cre to a dense granule protein (GRA) that is released into host cells only *<u>after</u>* invasion (Bougdour et al., 2013; Braun et al., 2013; Franco et al., 2016; Gay et al., 2016). Thus, GCre (GRA::Cre)-expressing parasites would not cause Cre-mediated recombination in the setting of aborted invasion (**Fig 2C**) and would identify only host cells that were or had previously been infected.

We engineered type II (Prugniaud) parasites to express an HA-tagged GCre (II-GCre). Using immunofluorescent assays and plaque assays, we determined that GCre was expressed, did not localize to the rhoptries, and that the expression of GCre did not affect overall parasite viability (**Fig S2**). To test the capability of II-GCre parasites to trigger Cre-mediated recombination, we infected fibroblast Cre reporter cells that express GFP only after Cre-mediated recombination (Koshy et al., 2010) with II-RCre parasites, II-GCre parasites, or parental type II parasites (no Cre expression). At 24 hours post-infection, we observed that both II-RCre parasites and II-GCre parasites caused host cell expression of GFP, while the parental strain did not (**Fig. S3A**). Compared to II-RCre parasites, II-GCre parasites showed a decreased efficiency of causing Cre-mediated recombination (>90% vs. 50%, **Fig. S3B**), which was expected because host cell-exported GRA proteins show decreased exportation when fused to ordered proteins (Bracha et al., 2018; Curt-Varesano et al., 2016; Franco et al., 2016).

Having confirmed that II-GCre parasites trigger Cre-mediated recombination, we next assessed the capability of II-GCre parasites to identify only infected cells. To address this concern, at 24 hours post-infection, we quantified the number of GFP^+^ Cre reporter fibroblasts that harbored parasites. With II-RCre parasites, ∼ 30% of GFP^+^ cells were actively infected, while with II-GCre parasites, ∼ 60% percent of GFP^+^ cells were infected (**Fig. S3B**). While the II-GCre parasites doubled the rate of infected GFP^+^ cells, ∼40% were uninfected. As this fibroblast Cre reporter cell line continues to divide after infection and Cre-mediated recombination (Koshy et al., 2012), we hypothesized that such division accounted for the uninfected GFP^+^ cells in II-GCre infected cultures. To test this possibility, we used cortical neuron cultures as neurons do not divide. In neuron cultures from Cre reporter mice, infection with II-RCre parasites resulted in 67% ± 1.82% of GFP^+^ neurons being actively infected, while infection with II-GCre parasites resulted in 98% ± 0.62% of GFP^+^ neurons being actively infected (**Fig 2 D, E**). Given that both RCre and GCre infected cultures showed a substantial increase in actively infected GFP^+^ cells when using non-dividing cells, these data suggest that in the fibroblast Cre reporter cell line, post-Cre-mediated recombination cell division accounts for ∼40% of the uninfected GFP^+^ cells. For the II-RCre parasites, the remaining ∼30% of uninfected GFP^+^ cells (and the ∼30% of uninfected GFP^+^ neurons) likely arise from aborted invasion (**Fig 2C**). For II-GCre parasites, that ∼100% of GFP^+^ neurons were infected confirms that II-GCre parasites trigger Cre-mediated recombination only after fully invading the host cell.

We next tested how IFN-γ pre-stimulation affected the rate of infected GFP^+^ neurons by stimulating Cre reporter neuron cultures with vehicle alone or IFN-γ (100 U/ml) for 24 hours prior to infection with II-GCre parasites. Consistent with previous results (**Fig 2E**), in the vehicle treated cultures, 97 ± 1.4% of GFP^+^ neurons harbored a parasite (**Fig. 2F, G**). In the setting of pre-treatment with IFN-γ, now only 78 ± 1.6% GFP^+^ neurons harbored parasites. The data suggest that the decrease in the rate of infection in IFN-γ stimulated neurons (**2A, B**) is primarily mediated by IFN-γ stimulated neurons clearing intracellular parasites, rather than IFN-γ stimulation leading to a decrease in parasite invasion.

### In IFN-γ stimulated murine neurons Irga6 loads onto the PVM in a Irgm1/3- dependent manner

As IFN-γ stimulated neurons up-regulate the IRG effectors (**Fig 1**) and a portion clear intracellular parasites (**Fig 2**), we next sought to determine if neurons use the IRG system to mediate IFN-γ-dependent killing of *T. gondii*. As the loading of IRGs on *T. gondii* PVM is indispensable for parasite clearance in IFN-γ stimulated non-neuronal cells (Y. O. Zhao et al., 2009) and as Irga6 is one of the effectors that loads onto the PV (Khaminets et al., 2010), we analyzed the percentage of PVs that were also Irga6^+^ in unstimulated and IFN-γ stimulated neurons infected with type III parasites. Consistent with findings in non-neuronal murine cells, we found that in unstimulated neurons, type III parasite PVs showed almost no Irga6 loading, while PVs in IFN-γ stimulated neurons showed an 8-fold increase in Irga6 loading (**Fig 3A, B**). To further confirm these findings, we also used a type III “IRG-resistant” strain (Cabral et al., 2016). This strain is engineered to express high levels of ROP18 (III+ROP18). ROP18 is a *T. gondii* kinase that phosphorylates Irga6 thereby preventing PV loading and effector oligomerization (Hermanns et al., 2016). The parental type III strain has minimal expression of ROP18 because of an insertion in the promoter region of the *rop18* gene; it is this lack of ROP18 expression that renders the parental strain susceptible to the IRGs (Saeij et al., 2006; Taylor et al., 2006). In cultures infected with III+ROP18 parasites, as expected, we now found almost no Irga6^+^ PVs even in the setting of IFN-γ stimulation (**Fig 3C, D**). Finally, to confirm that IRG-loading followed the same principles in neurons as in non-neuronal cells, we used neurons that lacked the regulatory IRG components (Irgm1 and Irgm3) (Collazo et al., 2001). Irgm1 and Irgm3 tether the effector components (e.g. Irga6) to the appropriate organelle until triggered to release the effectors onto the PV (Hunn et al., 2008) and thus are required for PV loading of Irga6 (Henry et al., 2009). In Irgm1/3 KO neurons, we found an increase in Irga6 dispersion throughout the cytosol in the setting of IFN-γ stimulation but no specific loading onto PVs regardless of IFN-γ stimulation or infecting strain (**Fig 3E-H**).

**Fig 3.**
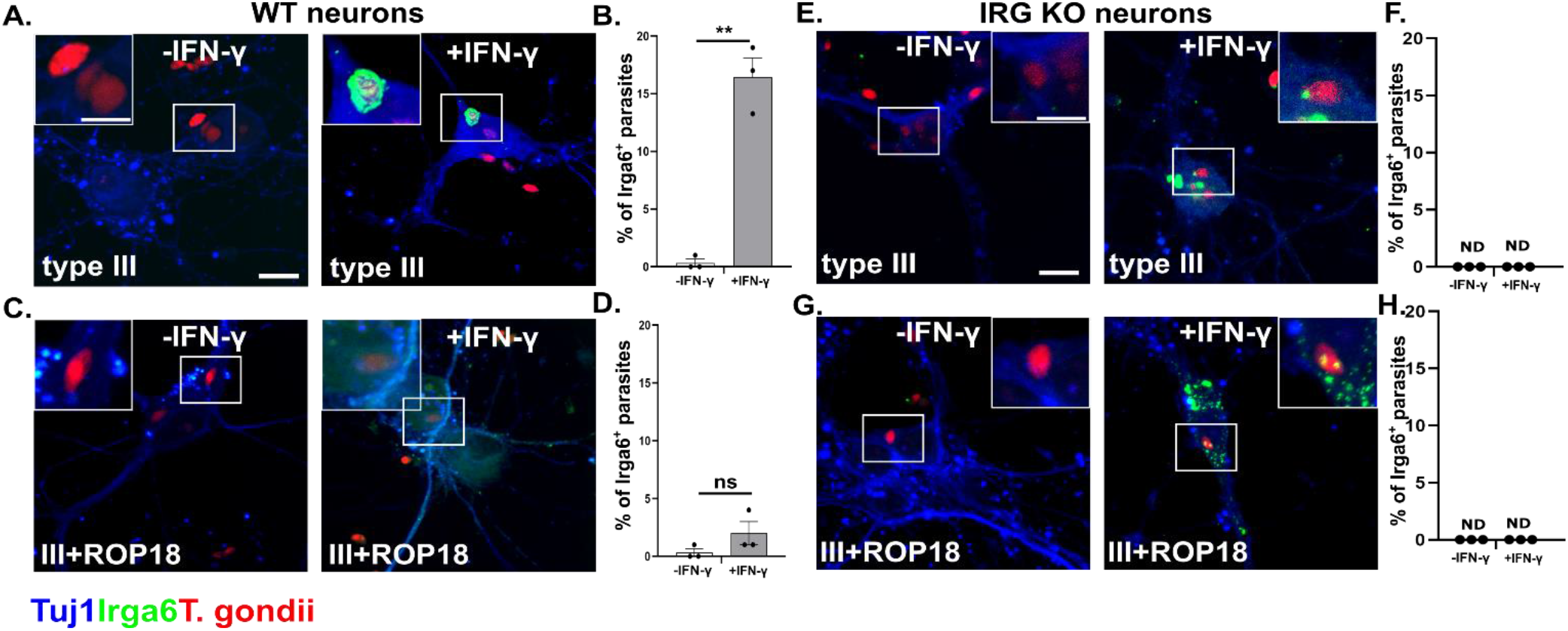
IFN-γ stimulated wild-type murine show increased loading of Irga6 onto the PVM of type III parasites. Primary neurons were cultured as in Fig 1, followed by 24 hours of IFN-γ (100U/ml) or vehicle stimulation, after which cultures were infected with listed *T. gondii* strains. At 12 hours post infection, cultures were fixed and stained with anti-Irga6 and anti-Tuj1 antibodies. The stained cultures were analyzed by confocal microscope. **A.** Representative images of stained cultures from wild-type (WT) mice infected with type III parasites and pre-stimulated with vehicle or IFN-γ. Blue = anti-Tuj1 antibodies, Green = anti-Irga6 antibodies, Red = mCherry expressing parasites. **B.** Quantification of the percentage of Irga6^+^ parasitophorous vacuoles (PVs) in the setting of vehicle or IFN-γ pre-stimulation. **C.** Representative images as in (**A**) except infected with III+ROP18 parasites. **D.** As in (**B**). **E.** Representative images as in (**A**) except using cultures from Irgm1/3 KO mice. Scale bar = 10 µm full image, 5 µm inset. **F.** As in (**B**). **G.** Representative images as in (**C**) except using IRG KO neurons. **H.** As in (**B**). (**B**, **D**, **F**, **H**) Bars- mean ± SEM, N = 100-200 PVs/experiment, 3 independent experiments. ns- not significant, **p ≤ 0.01, t-test with Welch’s correction.

Together, these data show that neuronal Irga6 loads onto the parasitophorous vacuole/PVM of intracellular IRG-sensitive parasites in the setting of IFN-γ pre-stimulation and when neurons have an intact IRG-system. These data strongly suggest that murine neurons use the IRG system for IFN-γ-dependent clearance/killing of intracellular parasites.

### IFN-γ stimulated murine neurons kill intracellular parasites using the IRG system

While the preceding data strongly suggests that murine neurons deploy the IRG-system to kill intracellular parasites in the settling of IFN-γ stimulation, they do not show that the IRG-system is essential for neuronal killing. To directly test this possibility, we took several approaches. First, we tested the ability of IFN-γ stimulated, wild type (WT) neurons to clear III+ROP18 parasites which show no Irga6^+^ loading even in the setting of IFN-stimulation (**Fig 3D**). Indeed, in murine neuronal cultures infected with III+ROP18 parasites, we found the same rate of infection over time, regardless of IFN-γ stimulation state (**Fig 4A**). Second, we infected Irgm1/3 KO neuronal cultures with parental (type III, IRG-sensitive) or III+ROP18 parasites (IRG-resistant) and with or without IFN-γ pre-stimulation. We again found the same rate of infection, regardless of what strain we utilized (IRG-sensitive or resistant) and IFN-γ pre-stimulation state (**Fig 4B**). As a final method of confirming that the lack of IRGs specifically affected intracellular parasites, we bred the Irgm1/3 KO mice to the Cre reporter mice to yield mice homozygous for the Cre reporter construct and that lack both Irgm1 and Irgm3 (Fig S4) and generated neuronal cultures from these Cre reporter Irgm1/3 KO mice. We then stimulated these neuronal cultures with vehicle or IFN-γ, followed by infection with II-GCre parasites. In these cultures, ∼98% (-IFN-γ: 98.12 ± 0.67%; + IFN-γ: 97.8% ± 0.53) of GFP^+^ neurons were infected, regardless of IFN-γ stimulation status (**Fig 4C, D**).

**Fig 4.**
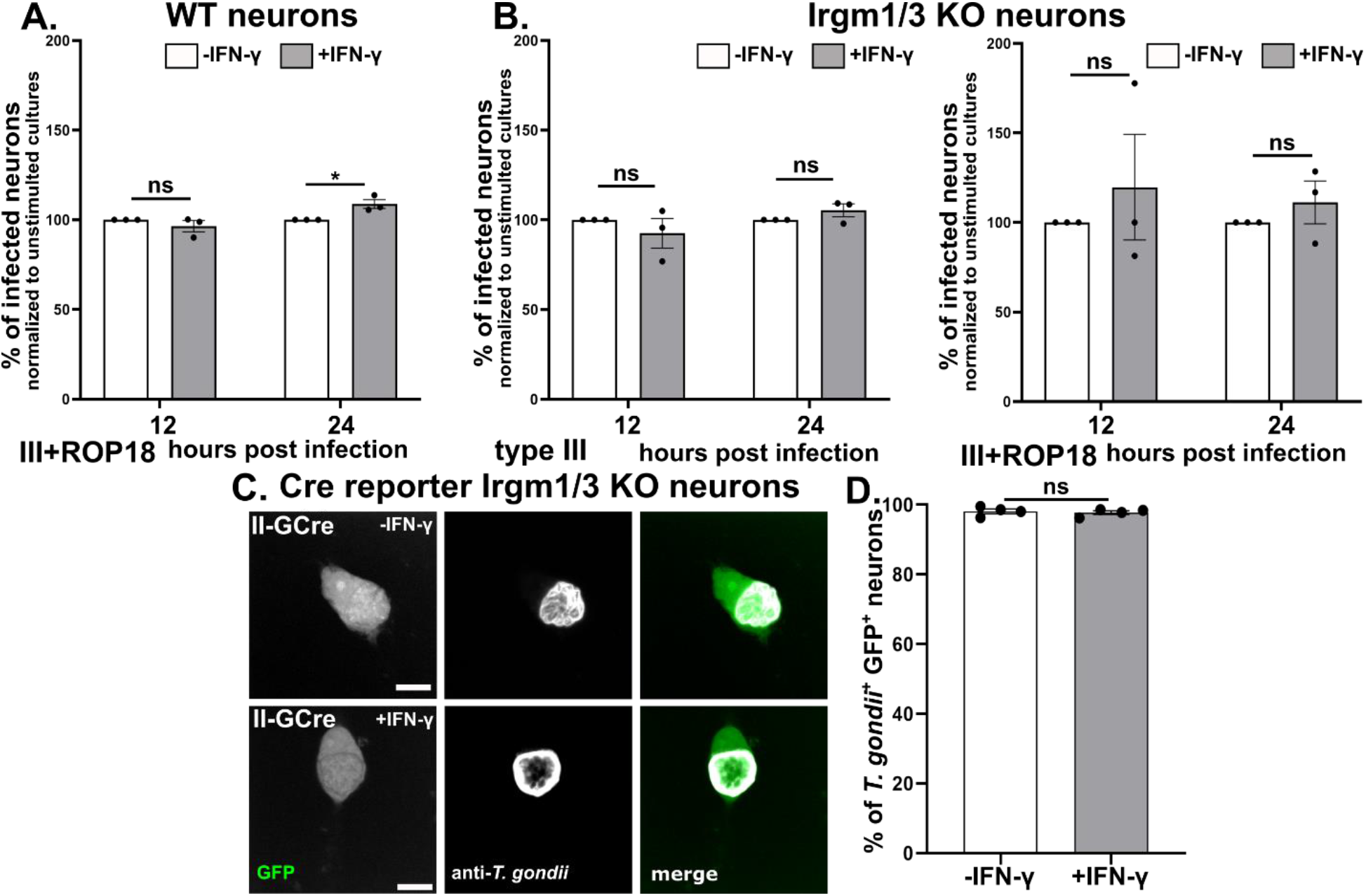
An intact IRG system is required for IFN-γ-dependent murine neurons clearance of intracellular parasites. Primary neurons from wild-type (Cre reporter), Irgm1/3 KO, or Cre reporter Irgm1/3 KO mice were cultured as in Fig 1 followed by stimulated with IFN-γ (100U/ml) or vehicle for 24hrs, after which the cultures were infected with listed *T. gondii* strain. **A.** Graph of the percentage of infected WT neurons at listed time points for III+ROP18 parasites, normalized to the unstimulated culture. **B.** As in (**A**) except using Irgm1/3 KO neurons and either type III parasites (*left graph*) or III+ROP18 parasites (*right graph*). (**A, B**) At the listed time points, cultures were fixed, stained, and analyzed as in Fig 2A**, B**. Bars mean ± SEM. N = 25-30 FOV analyzed/experiment (∼750-1000 neurons analyzed/experiment), 3 independent experiments. *p ≤ 0.05, ns = not significant, 2-way ANOVA with Sidak’s multiple comparisons. **C.** Representative images of GFP^+^ neurons from Cre Reporter Irgm1/3 KO mice infected with II-GCre parasites. *Merge image*: Green = GFP-expression in neurons, White = parasites stained with anti-*T. gondii* antibodies (a cocktail of anti-SAG1 and anti-SRS9 antibodies to capture both tachyzoites and bradyzoites.) Scale bars = 10 µm **D.** Graph of the percentage of infected GFP^+^ neurons. Bars, mean ± SEM. Eat dot represents the mean value of 1 experiment. N = N ≥ 200 GFP^+^ neurons analyzed/experiment, 4 independent experiments. ns = not significant, Welch’s t test. (**C, D**) Cultures were infected, fixed, stained, and analyzed as in Fig 2 **F, G**.

Collectively, these data definitively show that *in vitro* IFN-γ stimulated neurons kill intracellular IRG-sensitive parasites via the IRG system.

### Neurons clear intracellular parasites *in vivo*

Having shown that IFN-γ stimulated neurons clear intracellular parasites *in vitro*, we sought to determine if neurons cleared parasites *in vivo*. As the II-GCre parasites trigger Cre-mediated recombination only after full host cell invasion (**Fig 2C**), we reasoned that if we found GFP^+^, parasite^-^ neurons in Cre reporter mice infected with II-GCre parasites, these neurons must have cleared the invading parasite. To assess for GFP^+^ parasite- neurons *in vivo*, we created whole neuron reconstructions from 200 µm cleared brain sections stained with Hoechst from 21-day post-infection II-GCre infected mice (Cabral et al., 2020; Koshy and Cabral, 2014). As the PV excludes GFP expressed by the host cell, we looked for areas within the GFP^+^ neurons devoid of GFP (Fig 5A, B) and then confirmed that these areas contained parasites using Hoechst staining of the parasite DNA (Fig 5B). Out of 22 reconstructed GFP^+^ neurons, we found that 9 (∼40%) showed no evidence of persistent parasite infection (Fig 5C, D). These data highly suggest that *in vivo*, a percentage of murine neurons clear intracellular parasites.

**Fig 5.**
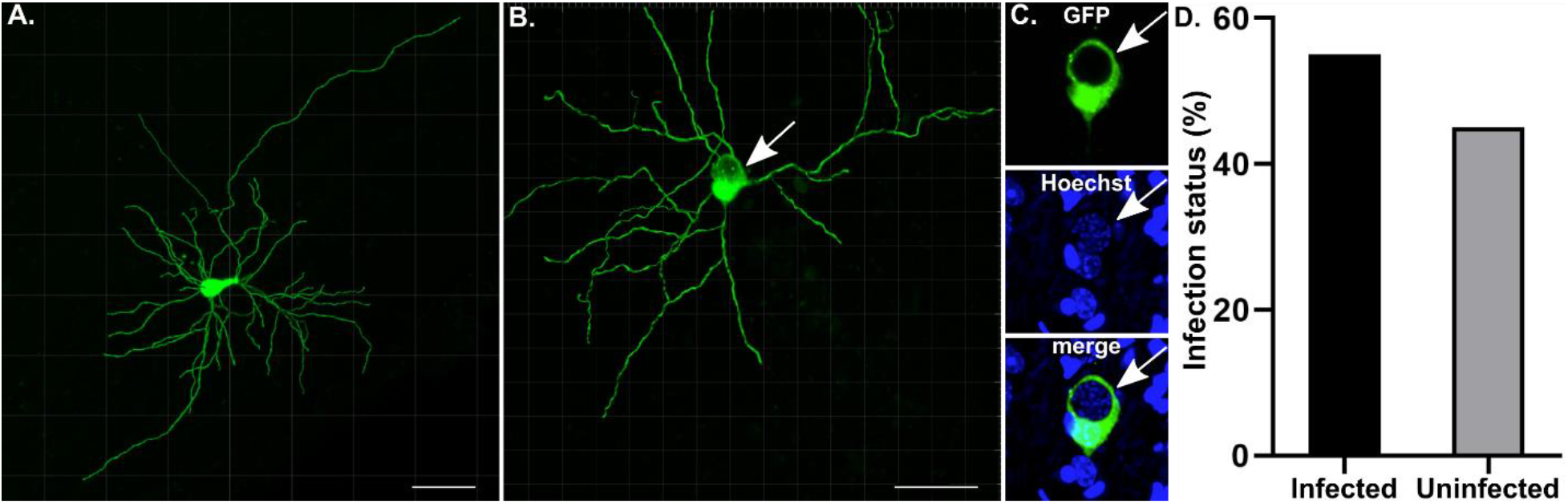
Forty percent of GFP^+^ neurons in mice infected with GCre parasites do not harbor parasites. Cre reporter mice were infected with GCre parasites. At 21 dpi, brains were harvested and sagittally sectioned into 200 µm thick sections. Thick sections were cleared and imaged at 40x on a confocal microscope. Neurons in resulting images were then rendered using Imaris software. **A.** A representative rendering of a GFP^+^ neuron in which no parasites were identified. **B.** As in (**A**) except now with a GFP^+^ neuron in which parasites were identified. White arrow shows parasites within neuron soma. **C.** Single plane of soma from (**B**) *top image*: GFP channel, *middle image*: blue channel (Hoechst), *bottom image*: merge. Note the GFP displacement, suggesting parasite presence within the neuron, which is then confirmed by visualization of parasite nuclei stained with hoecsht (blue). **D.** Graph of rendered GFP^+^ neurons containing parasites (infected) or not containing parasites (uninfected). N = 22 neurons from 4 mice.

### IFN-γ stimulated human neurons show resistance to *T. gondii* infection

Having shown that murine neurons clear a portion of intracellular parasites *in vitro* and *in vivo*, we sought to translate these findings to human neurons. While mice and murine cells are good models for human infection (both are naturally infected with *T. gondii*, have the CNS as a major organ of persistence, have neurons as the major host cell for cysts, and require IFN-γ and CD8 T cells to control toxoplasmosis), differences exist between the two. In the current context, the most relevant difference is that human cells lack the expansive range of Irgms that mice have and instead rely on alternative cell-specific mechanisms for IFN-γ-dependent control of *T. gondii* (Fisch et al., 2019). To determine how IFN-γ stimulation influenced control of *T. gondii* in human neurons, we derived human neurons (huSC neurons) from human neuroprogenitor cells reprogrammed from an embryonic stem cell line. After confirming that the huSC neurons expressed appropriate cortical neuronal markers (**Fig S5**), we used these huSC neurons for the *T. gondii* clearance assay. We pre-treated the huSC neurons with human IFN-γ for 24 hours, followed by infection with type II or III parasites. We then monitored the rate of neuron infection at 3, 12, and 24 hpi. At 3 hpi, regardless of infecting strain, we found equivalent neuron infection rates between unstimulated and IFN-γ stimulated cultures. At 12 and 24 hpi, IFN-γ stimulated cultures showed an ∼50% decrease in the number of infected neurons compared to unstimulated cultures (**Fig 6**).

**Fig 6.**
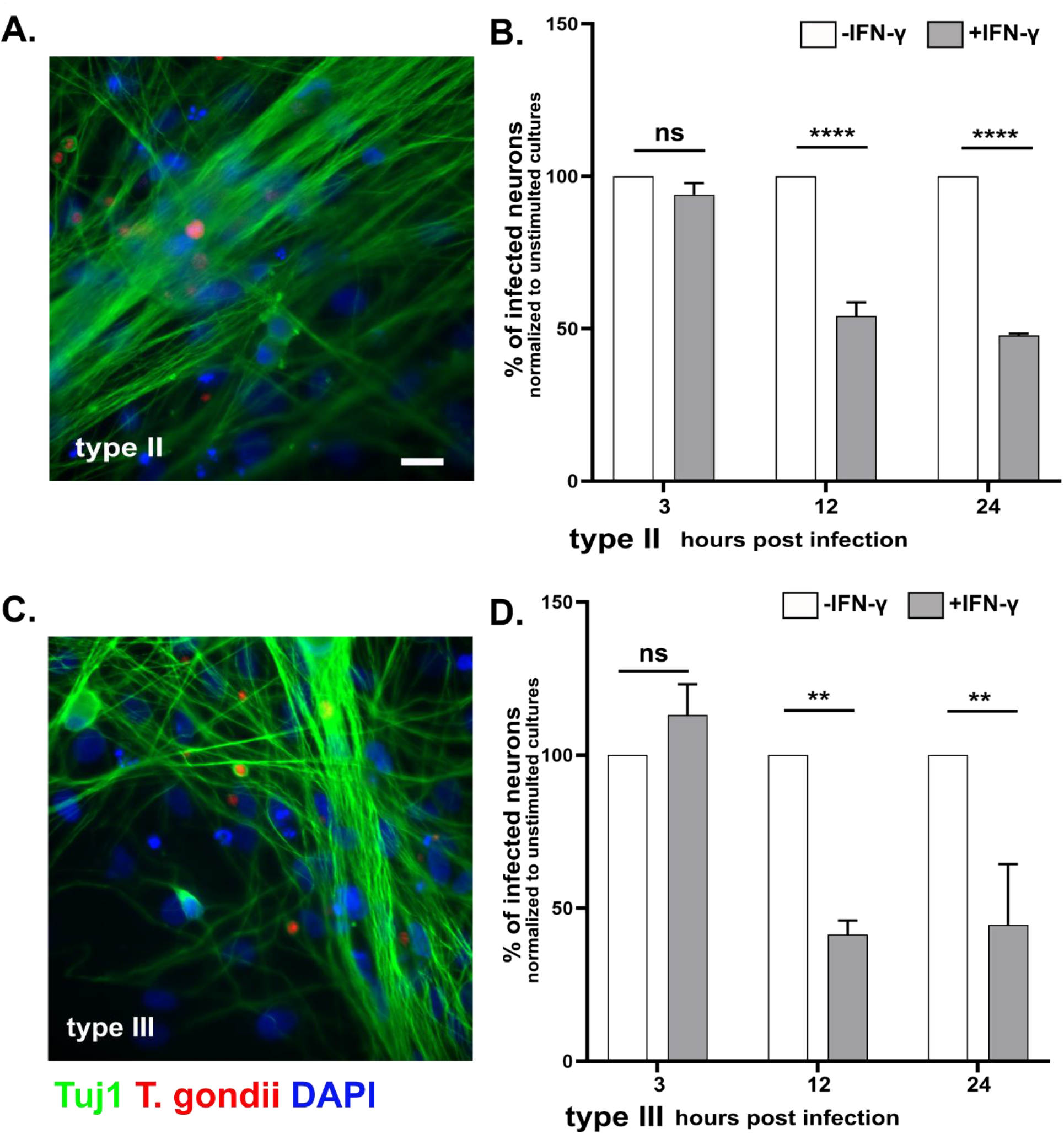
IFN-γ stimulation of human neurons leads to a decrease in the percentage infected with *T. gondii*. Human neurons were differentiated for 14 DIV from neural stem cells, after which they were stimulated with vehicle or IFN-γ (100 U/ml). At listed time points, cultures were fixed, stained with anti-Tuj1 antibodies and DAPI, and analyzed on an Operatta CLS high content analysis microscope. **A. C.** Representative images of human neurons infected with either type II or III parasites. Green = anti-Tuj1, Red = mCherry expressing parasites, Blue = DAPI. Scale bars = 20 µm **B.** Graph of the percentage of infected neurons at listed time points for type II parasites, normalized to the unstimulated culture. **D.** As in (**B**) except for type III parasites. Bars mean ± SD. N = 7-20 FOV (∼1000-1500 neurons/experiment), 2 independent differentiations. ****p<0.0001, **p<0.005 ns = not significant, 2-way ANOVA with Sidak’s multiple comparisons.

## Discussion

In this study, we sought to address the question: can neurons directly clear intracellular parasites? Using *in vitro* primary murine neurons from wild-type and transgenic mice in combination with IFN-γ pre-stimulation and transgenic parasites, this work shows that a portion of neurons can and do clear intracellular parasite in an IFN-γ-dependent, IRG- dependent manner. Using our new GCre parasites, which trigger host cell mediated recombination only after full invasion, we also show that ∼ 40% of neurons clear parasites *in vivo*. Finally, using huSC neurons, we translated our findings to human neurons, showing that IFN-γ pre-stimulation decreases the infection rate by ∼50%.

The data presented are the first to show that IFN-γ pre-stimulation enables human and murine neurons to partially resist infection by an intracellular eukaryotic pathogen. In murine neurons, we leveraged *T. gondii* biology to show that this resistance was secondary to clearance of intracellular parasites (**Fig 2A, B**), not simply an invasion defect (an important distinction in cultures where procedures such as aggressively washing off extracellular parasites cannot be done). In addition, our finding that IFN-γ stimulated murine neurons clear parasites in an IRG-dependent manner explains why a prior study found that IFN-γ stimulated primary murine neurons failed to clear intracellular parasites. The prior study, which was done at a time when neither the IRGs nor the parasite mechanisms to block the IRGs had been fully described, used a type I strain (RH) that we now know is IRG-resistant (Hermanns et al., 2016; Schluter et al., 2001). In human neurons, we currently cannot distinguish between an IFN-γ-dependent invasion defect or clearance of intracellular parasites or both. Though we cannot distinguish between these possibilities, the IFN-γ-dependent, anti-parasite effect appears to have a more robust effect on huSC neuron infection rates compared to murine neurons. What mechanisms underlie this impressive IFN-γ-dependent resistance to *T. gondii* infection will be the subject of future studies.

Though we have shown that IFN-γ-stimulated neurons can clear parasites in an IRG- dependent manner, our data also suggests that major cell-specific differences in IRG efficiency exist. We found that *in vitro*, ∼ 20-25% of intracellular parasites will be cleared by neurons in an IFN-γ, IRG-dependent manner, while other groups have shown that IFN-γ stimulated murine astrocytes, macrophages, and fibroblasts have higher rates of Irga6^+^ loading (50-75%) and clearance over a much shorter time (1-2 hours post-infection) (Khaminets et al., 2010; MacMicking, 2012; Martens et al., 2005). While some of the difference may be secondary to technical differences (e.g. use of antibody vs. transfection of Irga6-tagged with GFP, different MOIs), part of the difference is likely secondary to a blunted cell-intrinsic immune response from neurons (as suggested by the undetectable levels of baseline STAT1). Other, not mutually exclusive possibilities include that IRG-clearance differs between neuronal subcellular locations (i.e. it might be expensive to put IRG-machinery along the whole neuron) and/or that full neuron responses require direct interactions with other cell types such as astrocytes or T cells. Our *in vivo* data using GCre parasites are consistent with the possibility that other cell types influence neuronal clearance of intracellular parasites. *In vivo* we found that ∼ 40% of GCre-triggered GFP^+^ neurons do not harbor parasites (**Fig 5**). This rate of neuronal clearance is approximately double what we observed *in vitro*. In the *in vivo* setting, neurons are in constant communication with other cell types (e.g. astrocytes, microglia, infiltrating T cells), which may potentiate IRG-dependent clearance or initiate complementary methods for clearing intracellular parasites (e.g. CD40-dependent xenophagy (Andrade et al., 2006; Subauste, 2009)). Future studies will focus on defining these *in vivo* vs. *in vitro* differences.

In summary, our findings offer substantial evidence that IFN-γ pre-stimulation enables murine and human neurons to mount anti-parasitic defenses against *T. gondii*. While much work is left to be done to understand these anti-parasitic defenses, the work presented here suggests that *T. gondii’s* persistence in neurons is not simply a foregone conclusion.

## Acknowledgements

The authors would like to thank all members of the Koshy Lab for helpful discussions and critical review of the manuscript. We would like thank Greg Taylor (Duke University), for providing us with breeding pairs for *Irgm1/3^-/-^*, Jonathan Howard (Instituto Gulbenkian de Ciência), for providing us with anti-Irga6 antibodies. Finally, we thank Dr. Jared Churko (University of Arizona) and Dr. Rita Sattler (Barrow Neurological Institute) and their lab members for support with human neuron differentiations.

Funding was provided by the National Institutes of Health (NS095994 (AAK); AI147711 (JAK)) and the BIO5 Institute, University of Arizona (A.A.K). The funders had no role in study design, data collection and analysis, decision to publish, or preparation of the manuscript.

## Author Contributions

**Conceptualization**: Sambamurthy Chandrasekaran, Joshua Kochanowsky, Anita A. Koshy.

**Methodology**: Sambamurthy Chandrasekaran, Joshua Kochanowsky, Emily F. Merritt, Anita A. Koshy.

**Validation**: Sambamurthy Chandrasekaran, Joshua Kochanowsky, Emily F. Merritt

**Formal analysis**: Sambamurthy Chandrasekaran, Joshua Kochanowsky, Emily F. Merritt, Anita A. Koshy.

**Investigation**: Sambamurthy Chandrasekaran, Joshua Kochanowsky, Emily F. Merritt, Anita A. Koshy.

**Data curation**: Sambamurthy Chandrasekaran, Joshua Kochanowsky, Emily F. Merritt

**Writing – original draft**: Sambamurthy Chandrasekaran

Writing – review & editing: Anita A. Koshy

**Supervision**: Anita A. Koshy

**Funding acquisition**: Joshua A. Kochanowsky, Anita A. Koshy

## Declaration of Interests

The authors have declared that no competing interests exist.

## Materials and Methods

### Parasite maintenance

The parasite strains used in this study were maintained through serial passage in human foreskin fibroblasts (HFFs) using DMEM, supplemented with 10% fetal bovine serum (FBS), 2mM Glutagro and 100 IU/ml penicillin and 100 µg/ml streptomycin.

Except for the type II (Prugniaud) strain that expresses Gra16::Cre (II-GCre), the *T. gondii* strains used have been previously described (Cabral et al., 2016; Koshy et al., 2012). To engineer the II-GCre strain, type II parasites were electroporated with a plasmid encoding Cre recombinase fused to the dense granule protein, Gra16, and a separate drug-selectable marker (Donald et al., 1996). Single clones were selected by limiting dilution.

### Mice

All procedures and experiments were carried out in accordance with the Public Health Service Policy on Human Care and Use of Laboratory Animals and approved by the University of Arizona’s Institutional Animal Care and Use Committee (#12-391). All mice were bred and housed in specific-pathogen-free University of Arizona Animal Care facilities. Cre reporter mice (Madisen et al., 2010) (#007906) were originally purchased from Jackson Laboratories. Breeding pairs of *Irgm1/m3^-/-^* (Irgm1/3 KO) mice (Collazo et al., 2001) were generously provided by Greg Taylor (Duke University, Durham, NC).

### *In vivo* infection with GCre parasites

Cre reporter mice were intraperitonally infected with 10,000 or 20,000 freshly lysed Gra16-Cre (GCre) tachyzoites. Mice were anesthetized 21 days post infection, harvested brains were drop fixed in 4% PFA and stored overnight at 4^0^C before being transferred and stored in 30% sucrose until they were sectioned. *T. gondii* infected, sucrose embedded brains were sagittal sectioned to 200 µm on a vibratome and stored in cryoprotectant media (0.05 M sodium phosphate buffer containing 30% glycerol and 30% ethylene glycol). Sections were cleared using a modified PACT clearing protocol previously described (Cabral et al., 2020). In brief, sections were incubated overnight in a hydrogel monomer solution at 4 degrees Celsius and deoxygenated the next morning by bubbling nitrogen gas into sample vials. Samples were incubated at 42 degrees Celsius to initiate crosslinking of proteins, washed, and incubated in 8% sodium dodecyl sulfate (SDS) at 45 degrees Celsius to remove lipids. After multiple wash steps to remove SDS from the sections, nuclei were stained using hoechst. Samples were then washed and submerged in sorbitol refractive index matching solution (sRIMS) consisting of 70% Sorbitol and 0.01% NaN3 dissolved in 0.02M PB overnight before being mounted and imaged on a Zeiss NLO 880 confocal microscope (Imaging Core – Marley, University of Arizona). Brain sections were mounted for imaging on spacer slides in fresh sRIMS (Koshy and Cabral, 2014).

Images were obtained at 40x magnification to ensure visualization of parasite nuclei. We created z-stack tile scans of each neuron to ensure capturing as much of the neuron’s axon and processes as were available in each section. Stitched images were converted to Imaris files and imported into Bitplane Imaris software, where neuronal projections were rendered using the filaments tool.

### Primary murine neuron culturing

The primary neurons were cultured by methods described previously with minor modifications (Parker et al., 2018). The culturing plates were prepared by coating overnight with 0.001% poly-L-lysine (Sigma, Cat # P4707) solution for plastic surfaces and 100 µg/ml poly-L-lysine hydrobromide (Sigma, Cat # P6282) for glass surfaces.

Neurons were seeded in plating media at appropriate densities: 500,000 in 6-well plates for RNA and protein extraction, 100,000 in 24 well plates with coverslips for imaging and 20,000 in 96 well plates for counting. Four hours after plating, full volume media exchange to neurobasal media (Thermo Fisher, Cat # 21103049) was performed. On day in vitro (DIV) 4, neurons received a half volume media change of neurobasal media with 5 µM cytosine arabinoside to stop glial proliferation. One third media exchanges with neurobasal media occurred every 3-4 days thereafter. All the experiments were performed on 12 DIV neurons.

### IFN-γ stimulation and *T. gondii* infection

The primary neurons were pre-stimulated with 100 U/ml of mouse recombinant IFN-γ for 4h & 24h (RNA extraction), 24h & 48h (protein extraction), or 24h (*T. gondii* infections). Freshly syringe-lysed *T. gondii* parasites resuspended in neurobasal media were used to infect the primary neurons at MOI=4 (For 3 hpi), MOI=2 (For 12 hpi), MOI=1 (For 24 hpi) and MOI=0.2 (for 48 and 72 hpi) time points.

### Quantitative real time PCR

For quantification of the genes, RNA was extracted from 4h and 24h stimulated primary neurons using TRIzol reagent (Life technologies) and following the manufacturer’s protocol. 500 ng of total RNA was converted into first strand cDNA using a High-Capacity cDNA Reverse Transcription Kit (Applied Biosystems^TM^; Cat No: 4368814) and following the manufacturer’s instructions. Using the primer listed in Table 1, IFN-γ response and IRG pathway genes were amplified using SYBR green fluorescence detection with an Epppendorf Mastercycler ep realplex 2.2 system. GAPDH was used as a housekeeping gene to normalize DNA levels. Results were calculated using the 2^- ΔΔCT^ method (Livak and Schmittgen, 2001).

### Protein extraction and Western blotting

Primary neurons were either unstimulated or stimulated with IFN-γ for 24h and 48h, followed by total protein extraction as previously described (Cabral et al., 2017). Equal amounts of protein were subjected to SDS-PAGE, transferred to PVDF membrane and western blotting was done by standard methods. The blots were imaged using the Odyssey Infrared Imaging Systems (LI-COR Biosciences).

### Neuronal clearance assay

Primary neurons (wildtype or Irgm 1/3 KO) plated were plated on a poly-L-lysine coated 96-well plate (20,000/well) and were either unstimulated or pre-stimulated with IFN-γ for 24h prior to infection with *T. gondii* parasites. The cells were labeled with anti-NeuN (neuronal nuclei), DAPI (all nuclei) and parasites were mcherry positive, wells were imaged using an Operatta CLS high content analysis microscope (Functional Genomics Core, University of Arizona). Generated images were then analyzed using Image J software. The results were represented as percent decrease in the infected neurons in stimulated compared to the unstimulated group.

For assays involving GCre parasites, neurons from either wildtype (Cre reporter) or Cre reporter Irgm 1/3 KO mice were plated on poly-L-lysine coated 6 well plate (500,000/well). At the appropriate DIV, the neurons were either unstimulated or pre-stimulated with IFN-γ for 24h followed by infection with II-GCre parasites. At 72 hrs post-infection, the plates were processed as described above. These fixed and stained plates were then analyzed using an epifluorescent microscope (EVOS). The person analyzing the images was blinded to the IFN-γ stimulation and/or infecting parasite strain.

### Immunofluorescence assay

Cells were grown on poly-L-lysine-coated glass coverslips (described above) and were processed by methods as previously described (Parker et al., 2018).

### Antibodies

The following primary antibodies were used in the study: mouse anti-tubulin beta III isoform (Tuj1),clone TU20 (MAB1637, Millipore, 1:1000); rabbit anti- β3-Tubulin, D71G9 (similar to Tuj1) (5568S, CST, 1:1000); mouse anti-NeuN clone A60 (MAB377, Millipore, 1:1000); rabbit anti-Glial Fibrillary Acidic protein (GFAP) (Z0334, DAKO, 1:500); rabbit anti S100 (Z0311, DAKO, 1:500); rabbit anti-ALDH1L1 (Ab87117, Abcam, 1:500); chicken anti-Iba1 (Ab 139590, Abcam, 1:500), rabbit anti-STAT1 (Ab47425, Abcam; 1:500); mouse anti-pSTAT1 pY701 clone14/p-STAT1 (612132, BD Biosciences, 1:250); rabbit anti-pSTAT1 Tyr701, Clone 58D6 (9167, CST, 1:200); mouse anti-SAG1 DG52 (gift John Boothroyd, 1:10,000); mouse anti-SRS-9 (gift John Boothroyd, 1:10,000); mouse anti-Irga6 (1:1500), mouse anti-Irgb6 (1:250) (gift Jonathan Howard); rabbit ant- HA C29F4 (3724S, CST, 1:500); mouse anti-ROP2/3/4 (1:1000, gift John Boothroyd); DAPI (D3571, Thermo Fisher, 1:1000); Hoechst 33342 Trihydrochloride, Trihydrate (H3570, Thermo Fisher, 1:1000). The following species-appropriate secondary antibodies were used: Alexa Fluor 405 goat anti-rabbit IgG, Alexa Fluor 488 goat anti- mouse IgG, Alexa Fluor 568 goat anti-rabbit IgG, Alexa Fluor 647 goat anti-mouse IgG, Alexa Fluor 647 goat anti-chicken IgG, Alexa Fluor 647 goat anti-rabbit IgG, donkey anti-mouse IgG, DyLight 680 conjugate (1:10000), and donkey anti-rabbit IgG, DyLight 800 conjugate (1:10000). Unless otherwise noted, secondary antibodies were obtained from Life Technologies and used at a concentration of 1:500.

### Human stem cell derived neurons

H7 human embryonic Neural stem cells (NSCs) derived from the NIH-approved H7 embyronic stem cells (WiCell WA07) were purchased from the University of Arizona iPSC core (https://stemcells.arizona.edu/). The NSCs were expanded and differentiated into cortical layer neurons using a previously described protocol (Yan et al., 2013) with minor modifications. Briefly, NSCs were expanded on Matrigel^®^ Matrix (Corning^®^, #354277) coated plates using NSC expansion medium (NEM) (Thermofisher, Cat # A1647801). The media was changed every other day until NSCs reached confluence. The passaged NSCs were used at P2 for differentiation into cortical neurons by plated them on poly-L-ornithine (20ug/ml) (Sigma, Cat # P4957) and laminin (5ug/ml) (Thermo Fisher, Cat # 23017015) coated plates. For 14 days, the cells were differentiated into cortical neurons using neural differentiation medium (NDM) consisting of Neurobasal medium, 2mM L-Glutamine (Thermo Fisher, Cat # 25030024), 1% B-27 (Thermo Fisher, Cat # 17504044), 200µM L- Ascorbic acid (Sigma, Cat # A92902), 0.5mM c- AMP (Stem Cell Technologies Cat # 73886), 20ng/ml BDNF (Stem Cell Technologies Cat # 78005), 20ng/ml GDNF (Stem Cell Technologies Cat # 78058), 20ng/ml NT-3 (Stem Cell Technologies Cat # 78074), and Penicillin/Streptomycin (Thermo Fisher, Cat # 15140122) cocktail. The culture medium was exchanged with fresh NDM every 2-3 days.

### Human neuronal clearance assay

The clearance assay in human neurons was performed as described for primary pure murine neurons except that human IFN-γ (R&D Systems, Cat # 285-MI-100) was used for pre-stimulation.

### Statistical Analyses

Graphs were generated and statistical analyses were performed using Graphpad Prism 9.1.2 software. The specific test used (e.g., ANOVA vs. t-test) is noted in each figure.

---

**Fig S1.**
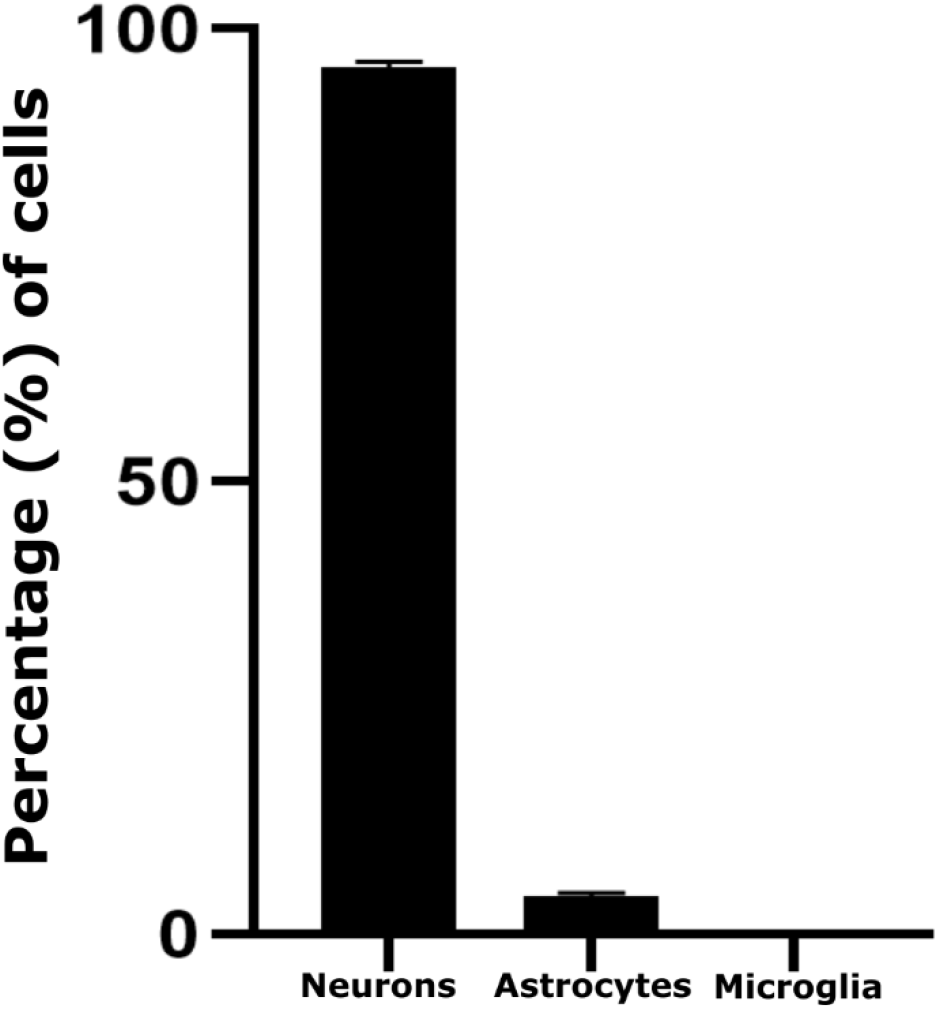
*In vitro* neuronal cultures are pure with little glial contamination. Quantification of the *in vitro* cultures after 12 DIV stained with anti-Tuj1 antibodies, (neuronal marker), an anti-astrocyte cocktail (anti-GFAP, anti-S100B, anti-ALDH1L1), and Iba1 (microglia marker). The numbers are presented as percentage of total cells counted (n= 100/experiment). Bars, mean ± SEM, N= 3 experiments.

**Fig S2.**
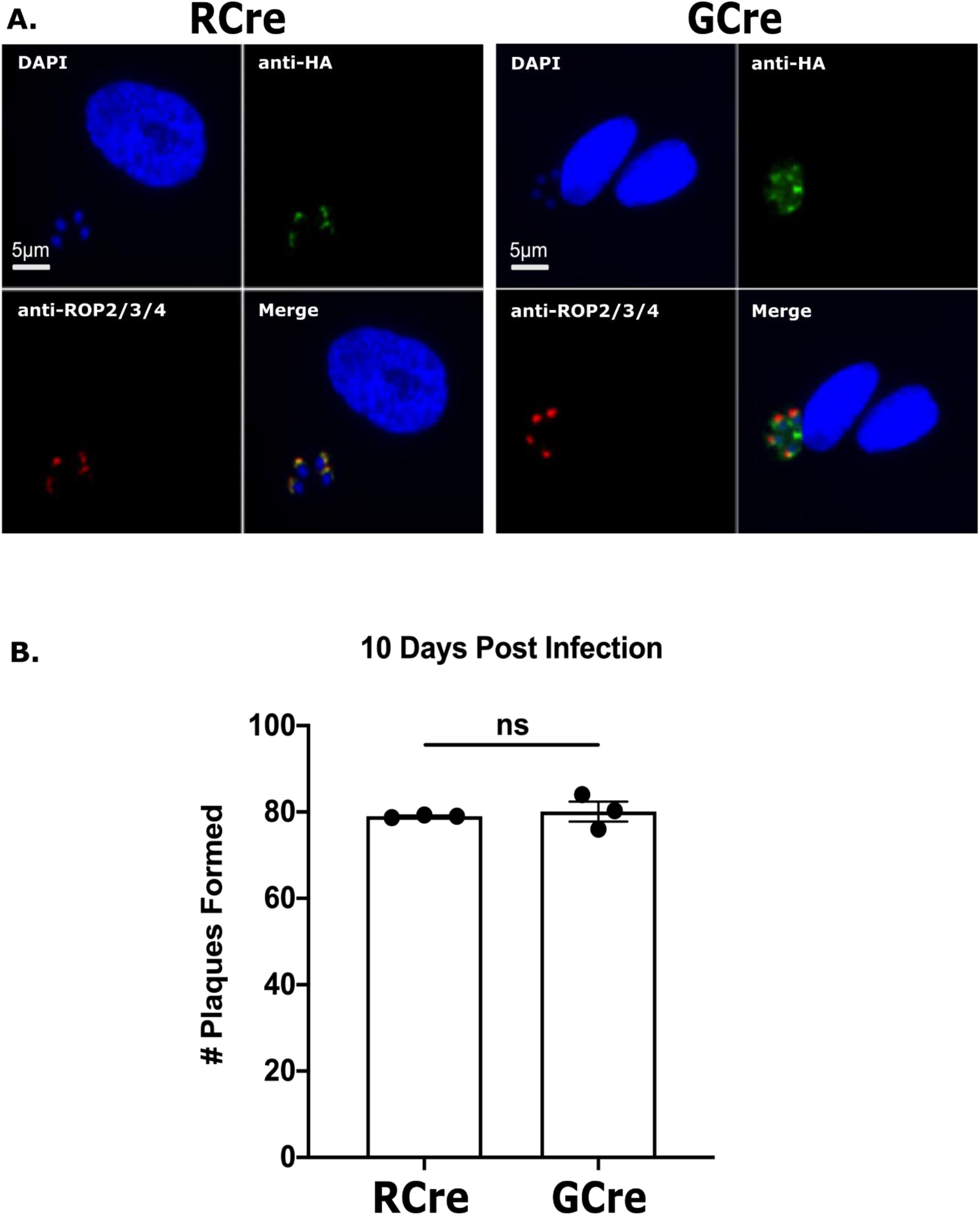
GCre is expressed and viable in *T. gondii*. **A.** Immunofluorescence for HA-tagged RCre and GCre fusion protein. HFFs were infected with identified strains (MOI=1) for 24 hours. Images depict listed *T. gondii*-Cre strain stained with anti-HA (green), anti-ROP2/3/4 (red), and DAPI (blue). **B.** Quantification of plaque assay at 10 dpi after infection with 200 parasites of the identified strains. Each dot = 1 experiment, N = 3 experiments. Bars, ± mean SEM. Welch’s t test. ns = not significant.

**Fig S3.**
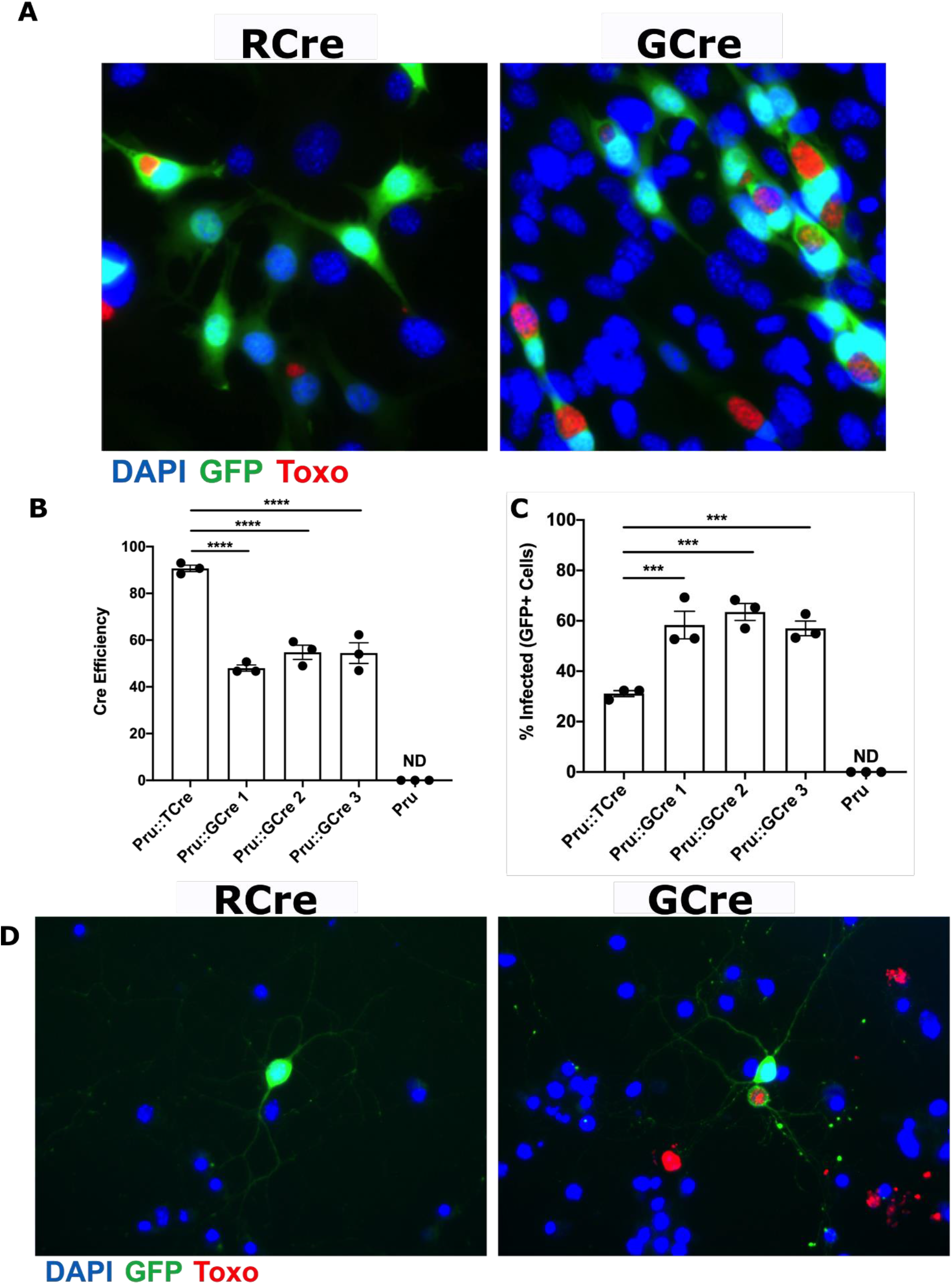
GCre is capable of Cre-mediated recombination in Cre-reporter fibroblasts and Cre-reporter neurons. **A.** Immunofluorescence for Cre-mediated recombination in Cre reporter fibroblasts infected with either RCre or GCre parasites. Cre reporter fibroblasts were infected with identified strains (MOI 1) for 24 hours and then fixed and stained. Images depict GFP (green), parasites (anti-SAG1; red), and nuclei (DAPI, blue). **B.** Quantification of Cre-mediated recombination efficiency. Cre reporter fibroblasts were infected as in (**A**) and efficiency scores were quantified by dividing the number of GFP^+^ cells by the number of infected cells multiplied by 100. **C.** The percentage of GFP^+^ cells that actively harbored a parasite was quantified by dividing the number of infected GFP+ cells by the total number of GFP^+^ cells multiplied by 100. **D.** Immunofluorescence for Cre-mediated recombination in Cre reporter neurons infected with either RCre or GCre parasites. Cre reporter neurons were infected with identified strains (MOI=0.1) for 72 hours and then fixed and stained. Images depict GFP^+^ neurons (green), parasites (anti-SRS9 and anti-SAG1; red), and nuclei (DAPI, blue). **B** and **C** Values, mean ± SEM. Each dot represents the mean value of 1 experiment. N = 3 experiments. One-way ANOVA with Dunnett’s multiple comparison test. ***p<0.0005 and ****p<0.0001. ND = Not Detected.

**Fig S4.**
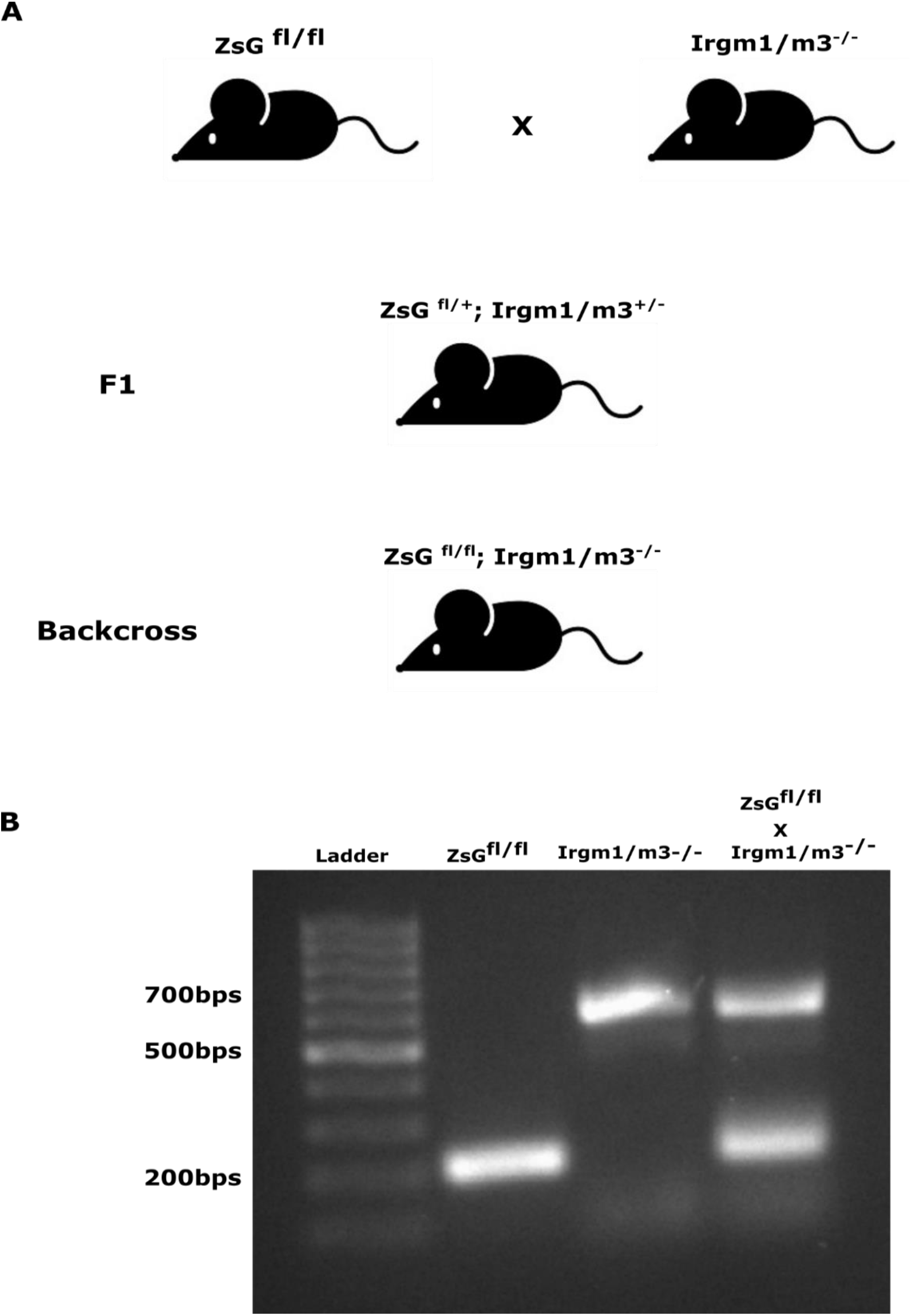
Generation of Cre reporter Irgm1/m3 knockout mice. **A.** Schematic representation of breeding Cre reporter to Irgm1/m3 knockout mice. **B.** Agarose gel representing the genotyping of the transgenic lines.

**Fig S5.**
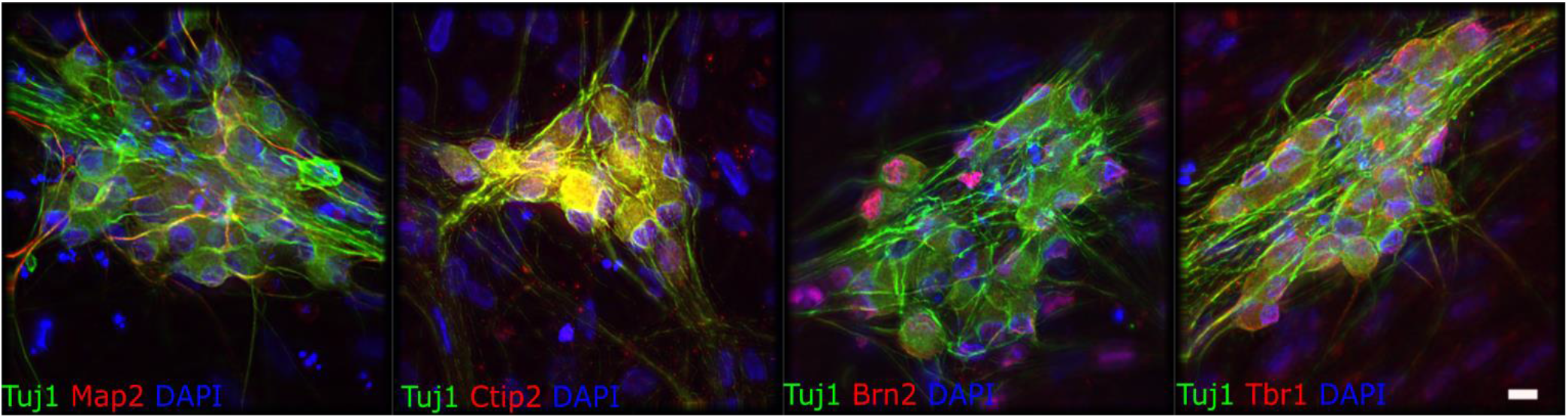
Characterization of human neurons for cortical layer markers. Human stem cell derived neurons were grown and differentiated for 14 days, after which cultures were fixed and stained with DAPI and anti-Tuj1 antibodies (pan neuronal marker) and one of the following: anti-Map2 antibodies (mature neuronal marker), anti- Ctip2 antibodies (cortical layer V, VI), anti-Brn2 antibodies (cortical layer II-V), or anti- Tbr1 antibodies (cortical layer I, V, VI). Scale bar = 25µm

## Notes

### Competing Interest Statement

The authors have declared no competing interest.

### Summary of Updates

Figure 2, 4 and 6 updated; Author list updated.

